# Challenges in the Biotechnological Implementation of Oral RNA Interference as an Antiviral Strategy in *Aedes aegypti*

**DOI:** 10.1101/2023.11.05.565667

**Authors:** Ottavia Romoli, Annabelle Henrion-Lacritick, Hervé Blanc, Lionel Frangeul, Maria-Carla Saleh

## Abstract

Mosquitoes, particularly *Aedes aegypti*, are critical vectors for globally significant pathogenic viruses. This study examines the limitations of oral RNA interference (RNAi) as a strategy to disrupt viral transmission by *Ae. aegypti*. We hypothesized that double-stranded RNA (dsRNA) targeting the Zika virus (ZIKV) or chikungunya virus (CHIKV) genomes produced by engineered bacterial symbionts could trigger an antiviral response. Mosquitoes mono-colonized with *Escherichia coli* producing dsZIK or dsCHIK did not display reduced viral titers following exposure to virus-contaminated bloodmeals and failed to generate dsZIK- or dsCHIK-derived small interfering RNAs. To address potential limitations of bacterial dsRNA release, we explored dsRNA inoculation via feeding and injection. While viral replication was impeded in mosquitoes injected with dsZIK or dsCHIK, no antiviral effect was observed in dsRNA-fed mosquitoes. These findings highlight complexities of implementing oral RNAi as an antiviral strategy in *Ae. aegypti* and warrant further exploration of local and systemic RNAi mechanisms.

## Introduction

Mosquitoes are vectors for several arthropod-borne viruses (or arboviruses), which are responsible for multiple outbreaks worldwide. Over the past decade, the emergence or re-emergence of viruses such as dengue (DENV), Zika (ZIKV), or chikungunya (CHIKV) viruses, together with the current lack of vaccines for most of these viruses and the emergence of insecticide resistant mosquito populations has necessitated the development of new strategies to control the spread of these pathogens and their vectors.

Mosquito vector competence (*i.e.*, the ability of a mosquito to acquire and transmit a virus) is dependent on efficient viral replication and dissemination from the gut to the salivary glands, where the virus must enter the saliva to be transmitted to the next vertebrate host during subsequent blood-feeding. One of the mechanisms that mosquitoes use to control viral replication is RNA interference (RNAi), which is triggered by viral double-stranded RNA (dsRNA) intermediates produced during RNA virus replication. The RNase Dicer-2 binds dsRNAs and processes them into 21 nucleotide (nt) small-interfering RNAs (siRNAs)^1^. These are used by Argonaute-2 as sequence-specific guides to target complementary viral RNAs, thereby hindering viral replication^2–5^. Key studies in *Aedes aegypti* mosquitoes showed that a protective RNAi-mediated antiviral response could be triggered by exposure to dsRNA molecules corresponding to portions of the DENV or ZIKV genomes either by transgenic production of dsDENV or by injection of dsZIK^6,7^, validating the potential use of exogenous dsRNA molecules to limit arbovirus infection.

During viral infection through a bloodmeal, the mosquito midgut represents the first mechanical and immunological barrier encountered by the virus and crossing this barrier is critical to establish an infection in the mosquito. Thus, we reasoned that blocking viral replication in the midgut could be an effective way to limit viral transmission and we hypothesized that bacterial symbionts in the mosquito gut could be used as a platform to produce and deliver dsRNA to mosquitoes and stimulate a prophylactic RNAi-based antiviral response. Indeed, a recent study in honeybees showed the possibility of reducing viral replication through colonization with an antiviral dsRNA-producing symbiont^8^. When it comes to mosquitoes, numerous studies have demonstrated the potential for gene silencing by introducing dsRNA through feeding or using microorganisms that produce dsRNA^9–14^. While a handful of studies have pointed out challenges like low efficiency and variability in oral RNA interference in mosquitoes^15,16^, it’s noteworthy that none of these investigations have explored the concept of microbiota-mediated oral delivery of antiviral dsRNA.

Paratransgenesis is the genetic modification of a symbiotic microorganism that is associated with a disease vector with the aim of disrupting the transmission of pathogens carried by the vector. This approach offers the advantage of potential self-sustainability, as the targeted microorganism is typically selected to undergo effective horizontal (between individuals) and vertical (from parents to progeny) transmission^17^. Various paratransgenic methods have been proposed to reduce *Plasmodium* transmission by *Anopheles* mosquitoes, including the use symbiotic bacteria engineered to produce anti-*Plasmodium* molecules or mosquito immune factors^18–20^. Additionally, other paratransgenic strategies have focused on interfering with mosquito development or fertility^10,12,21^. However, there are currently no developed strategies to impede virus transmission by *Aedes* mosquitoes through manipulation of their microbiota.

Here, we investigated the potential targeted induction of antiviral RNAi against ZIKV and CHIKV in *Ae. aegypti* mosquitoes through colonization with bacteria capable of producing antiviral dsRNA. We used *Escherichia coli* as a proof-of-concept and found that mosquitoes colonized by dsZIK- and dsCHIKV-expressing bacteria did not exhibit reduced viral titers when infected with the corresponding virus through a bloodmeal. Additionally, there was no discernible production of siRNAs induced by the ingestion of bacteria-produced dsRNA. When we fed mosquitoes with naked dsRNA, we did not observe enhanced antiviral immunity, likely due to the absence of an effective antiviral RNAi-based response in the midgut and insufficient activation of systemic RNAi.

## Results

### Colonization of mosquitoes with dsRNA-producing Escherichia coli

To generate *E. coli* strains expressing antiviral dsRNA, we took advantage of the HT115 strain, which has historically been used in *Caenorhabditis elegans* to induce RNAi. This strain is suitable for dsRNA production as it possesses an isopropylthio-β-galactoside (IPTG)-inducible *T7 polymerase* gene and has a mutation in the *rnc* allele encoding RNase III, resulting in reduced dsRNA degradation^22^. This allows the synthesis of the target sequence by transcription from a DNA template containing the T7 promoter. To induce the production of dsRNA for ZIKV or CHIKV, we cloned the target viral sequences between two directionally opposed T7 promoters and terminators in the T444T plasmid to produce dsRNA by convergent transcription^23^ (**Figure 1A, Supplementary Figure 1A**). To test whether a unidirectionally-transcribed RNA with a hairpin secondary structure would result in a more stable and effective dsRNA production, an inverted repeat of the CHIKV target sequence was cloned in a pUC18 plasmid downstream from a T7 promoter. The inverted repeat consisted of the CHIKV target sequence in both polarities separated by a non-viral spacer sequence (**Supplementary Figure 1B**). The target viral sequences were selected for dsRNA production based on the following criteria: (i) we excluded sequences that could produce 21mers with high similarity to any sequence in the *Ae. aegypti* genome in order to avoid potential interference of endogenous gene expression; (ii) we targeted regions corresponding to only partial coding regions of viral structural genes to prevent translation of the target sequences into functional viral proteins; (iii) we avoided sequences corresponding to noncoding subgenomic flaviviral RNA or viral proteins shown to repress mosquito immunity^24–26^(**Figure 1B, Supplementary Figure 1C**). A T444T plasmid containing a portion of the GFP coding sequence instead of a target viral sequence was produced as a control.

**Figure 1.**
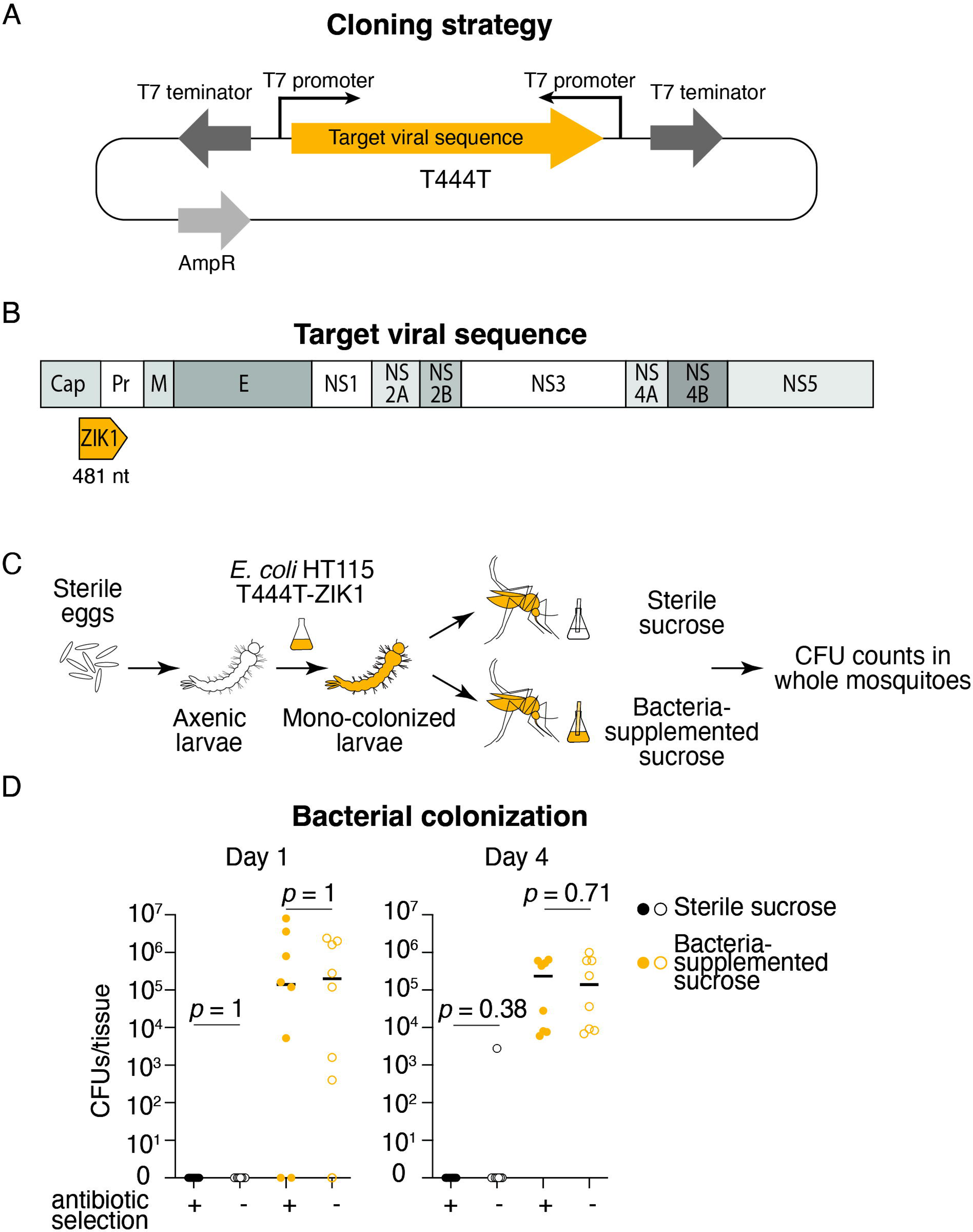
Colonization of mosquitoes with dsRNA producing *E. coli*. (**A**) Representation of the T444T plasmid. The target viral sequence was cloned between two directionally opposed T7 promoters to allow convergent transcription of a sense and antisense RNAs that form dsRNA upon annealing. (**B**) Position of the target viral sequence on the ZIKV genome. The selected sequence is 481 nt and spans the coding sequences for the Capsid (Cap) and the Pre-membrane (Pr) protein coding genes. The other components of the ZIKV genome are: M (Membrane), E (Envelope) NS1-5 (Non-Structural proteins). (**C**) Experimental set up for mosquito mono-colonization with *E. coli* HT115. Sterile eggs were obtained by subsequent washes in 70 % ethanol and 1 % bleach. An overnight culture of *E. coli* HT115 carrying the T444T-ZIK1 plasmid was added to axenic larvae hatched from sterile eggs to obtain mono-colonized larvae. Adults emerging from mono-colonized larvae were provided with either sterile sucrose or sucrose supplemented with *E. coli* HT115. Colony forming units (CFUs) were determined in whole adults to evaluate the efficiency of bacterial colonization. (**D**) Bacterial loads (mean) and plasmid stability *in vivo*. CFUs were determine one and four days-post-emergence in adult mosquitoes mono-colonized with *E. coli* HT115-T444T-ZIK1 at the larval stage, fed with either sterile (black) or bacterial supplemented (yellow) sucrose. To determine plasmid stability in bacteria during mosquito colonization, CFUs were determined on LB plates supplemented (full dots) or not (empty dots) with ampicillin. Bacterial loads were compared using a Wilcoxon test.

To confirm the production of dsRNA by bacteria, RNA was extracted from overnight cultures of *E. coli* HT115-T444T-ZIK1 induced or not with IPTG for 24 h. Northern blot with ZIK1 sense and antisense probes was performed on bacterial total RNA and on dsZIK1 synthesized *in vitro* as a positive control. The agarose gel showed a major 480 nt band for *in vitro* synthesized dsZIK1, although a higher ∼900 nt band was visible (**Supplementary Figure 2A**). In the corresponding northern blot, ZIK1 sense and antisense probes bound to both 480 and ∼900 nt bands (**Supplementary Figure 2B**), suggesting that the larger band is composed by dsRNA multimers or higher order assemblies. These molecules are commonly found in both *in vitro* and *in vivo* synthesized dsRNA with unknown effect on RNAi activity^27,28^. A signal at ∼350 nt was detected by sense and antisense probes in uninduced bacteria (**Supplementary Figure 2B**). This might represent a truncated RNA transcribed by the T7 RNA Polymerase, whose transcription is directed by *lac* operon control elements. This system is commonly known to be leaky and to show basal transcription in absence of IPTG induction, especially in stationary phase cultures^29,30^. Bacteria induced with IPTG showed a major ∼650 nt band in the agarose gel (**Supplementary Figure 2A**), which was recognized by both ZIK1 probes together with other transcription products spanning from ∼250 to ∼900 nt (**Supplementary Figure 2B**). The major band at ∼650 nt represents the main ZIK1 transcript from the T444T-ZIK1 plasmid, which is 186 nt longer than the *in vitro* transcript due to additional plasmid sequences flanking the *Not*I and *Age*I cloning sites. The ZIK1 sense probe showed high specificity for the target ZIKV sequence, while no significant signal was visible in a northern blot of total RNA extracted from overnight cultures of *E. coli* HT115-T444T-CHIK1 (**Supplementary Figure 2C-D**). Altogether the northern blot analysis revealed abundant production of dsRNA by the modified bacteria.

To test the mosquito colonization properties of *E. coli* HT115, the transstadial transmissibility of *E. coli* HT115 (from larvae to adults), and the stability of the T444T plasmid *in vivo* in the absence of antibiotic selection, we produced *Ae. aegypti* larvae mono-colonized by *E. coli* HT115-T444T-ZIK1 (**Figure 1C**). Mono-colonization at the larval stage was used to maximize the exposure of mosquitoes to dsRNA. *E. coli* HT115 was added to axenic larvae obtained from surface sterilized eggs^31^. The HT115 strain was able to support standard larval development, as previously described for other *E. coli* strains^31,32^. To assess the transstadial transmission of *E. coli* HT115, we measured bacterial loads in adult mosquitoes originating from mono-colonized larvae kept in non-sterile conditions after adult emergence. When mosquitoes were provided only with sterile sucrose, *E. coli* HT115 carrying the T444T-ZIK1 plasmid could not be isolated from whole mosquitoes collected one or four days post-emergence, showing the low transstadial transmissibility of this strain (**Figure 1D**). When mosquitoes were provided with a sucrose solution contaminated with *E. coli* HT115 carrying the T444T-ZIK1 plasmid, bacteria could be isolated from whole mosquitoes at both tested time points, with 100 % of mosquitoes colonized four days post-emergence. Bacterial loads were comparable when measured on antibiotic-free plates or plates containing ampicillin to select for bacteria carrying the T444T-ZIK1 plasmid (Wilcoxon test: sterile sugar day 1, *p* = NA; sterile sugar day 4, *p* = 0.38; bacteria supplemented day 1, *p* = 1; bacteria supplemented day 4, *p* = 0.71), indicating that *E. coli* HT115 could stably colonize adult mosquitoes, it represented most of the bacteria present at the adult stage, and it could maintain the plasmid *in vivo* for at least four days.

### Mosquitoes colonized with dsRNA-producing bacteria do not show reduced viral titers or production of dsRNA-specific siRNAs

To assess the effect of the bacterial-produced dsRNA on viral infections, we exposed mosquitoes to a ZIKV-or CHIKV-containing infectious bloodmeal. We reasoned that mosquitoes mono-colonized at the larval stage with the dsRNA-producing bacteria would be the best model to work with, as they were exposed to the dsRNA during their entire development. We produced larvae mono-colonized with *E. coli* HT115 carrying the T444T-GFP, T444T-ZIK1, T444T-CHIK1, or pUC18-CHIK1 plasmids and we provided the same bacteria to the adult mosquitoes that developed from these larvae (**Figure 2A, Supplementary Figure 1D**). To activate transcription of the dsRNA from the T7 promoter(s), IPTG was added to the bacterial culture medium, the larval water, and the sucrose solution provided to adults. Mosquitoes were offered bacteria-contaminated sucrose from their emergence onward. Four-to five-day-old mosquitoes were blood-fed with a ZIKV-or CHIKV-containing infectious bloodmeal and provided with bacteria-contaminated sucrose after the bloodmeal. For ZIKV, viral titers were measured in the midguts, carcasses, and heads of mosquitoes at six and ten days post-infection to assess infection and dissemination rates. We could not detect a significant difference in ZIKV titers for mosquitoes colonized by dsZIK1-producing *E. coli* compared to mosquitoes colonized by dsGFP-producing *E. coli* in any tissue at any time-point (**Figure 2B, C,** Wilcoxon test on log transformed viral titers: day 6 midguts *p* = 0.54, carcasses *p* = 0.72, heads *p* = 0.87; day 10 midguts *p* = 0.09, carcasses *p* = 0.29, heads *p* = 0.16). The prevalence of infection was also similar between the two conditions (Wilcoxon test: day 6 midguts *p* = NA, carcasses *p* = 0.71, heads *p* = 0.63; day 10 midguts *p* = 0.55, carcasses *p* = 0.35, heads *p* = 0.29). These data indicate that bacteria-produced dsRNA did not induce a protective RNAi-based antiviral response.

**Figure 2.**
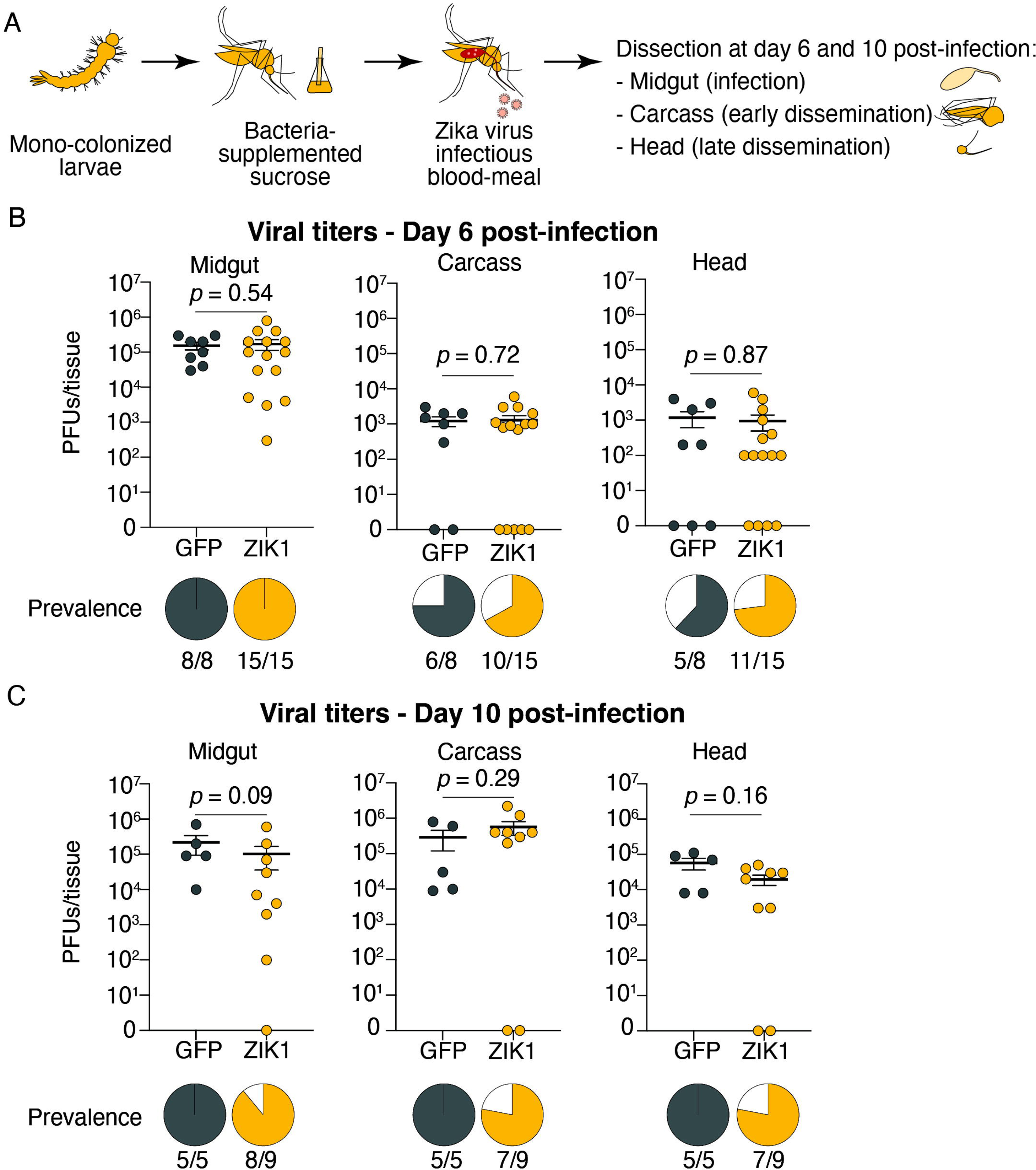
Mosquitoes colonized with dsZIK-producing bacteria do not show reduced viral titers. (**A**) Experimental protocol for ZIKV infection of mosquitoes colonized by dsRNA-producing *E. coli*. Mosquitoes originating from mono-colonized larvae were kept on bacteria-supplemented sucrose for four days. After one day of starvation, they were offered a ZIKV infectious bloodmeal and subsequently kept with bacteria-supplemented sucrose until sampling at six and ten days post-infection. At each time point midguts, carcasses, and heads of individual mosquitoes were collected. (**B-C**) Viral titers (mean ± SEM) in midguts, carcasses, and heads of individual mosquitoes colonized by *E. coli* HT115 expressing dsGFP (grey) or dsZIK1 (yellow) and collected six (**B**) or ten (**C**) days post-infection. Each plot shows the log of plaque forming units (PFUs) measured in individual mosquitoes in each tissue. Pie charts show the proportion of infected (grey/yellow) and uninfected (white) mosquitoes. Numbers below pie charts represent the number of infected over uninfected mosquitoes per tissue/time-point. Viral loads were compared using a Wilcoxon test.

For CHIKV, viral titers were measured in the bodies and heads of mosquitoes at two and five days post-infection. The results indicated that CHIKV titers were not specifically reduced by the exposure of mosquitoes to dsCHIKV. Contrary to our hypothesis that dsRNA-producing bacteria could induce a sequence-specific prophylactic antiviral response, CHIKV titers were higher in the bodies and heads of mosquitoes colonized with bacteria carrying the T444T-CHIK1 or pUC18-CHIK1 plasmids than those of mosquitoes colonized with bacteria carrying the T444T-GFP plasmid at day two post-infection, while viral loads were similar in mosquitoes colonized with the three types of bacteria at day five post-infection (**Supplementary Figure 1E-F**, Kruskal-Wallis test on log transformed viral titers: day 2 body *p* < 0.001; day 2 head p = 0.004; day 5 body *p* = 0.3, ns; day 5 head *p* = 0.8, ns; Kruskal-Wallis test on prevalence day 2 body *p* = NA; day 2 head p = 0.003; day 5 body *p* = NA; day 5 head *p* = 0.5, ns; see Supplementary Table 2 for individual comparisons). Taken together, these results suggest that colonization of mosquitoes with dsRNA-producing bacteria did not trigger an efficient antiviral response against flaviviruses or alphaviruses.

To verify whether the mosquito RNAi machinery could produce siRNAs from the dsRNA produced by bacteria, we sequenced small RNAs from adult mosquitoes mono-colonized for two days with *E. coli* HT115 carrying the T444T-ZIK1 plasmid (**Figure 3A**). Reads mapping to the ZIKV target sequence on both positive and negative strands were obtained, with an enrichment of positive strand-reads (**Figure 3B**). This further confirmed that dsRNA is effectively transcribed *in vivo* by *E. coli* HT115. Most of the reads mapping to the target viral sequence fell within the expected range of 20 to 28 nt, but no enrichment at 21 nt was observed, suggesting that siRNAs were not produced from the bacterial-produced dsRNA (**Figure 3C**). As our mosquito colony is persistently infected with the insect specific virus Phasi-Charoen Like Virus (PCLV), we used this natural infection as a quality control for our small RNA sequencing. We observed that 21 nt reads were significantly enriched among all small RNA reads from 18 to 33 nt, suggesting that PCLV-specific siRNAs were produced in these mosquitoes and were detected by our small RNA sequencing pipeline (**Supplementary Figure 3A-F**).

**Figure 3.**
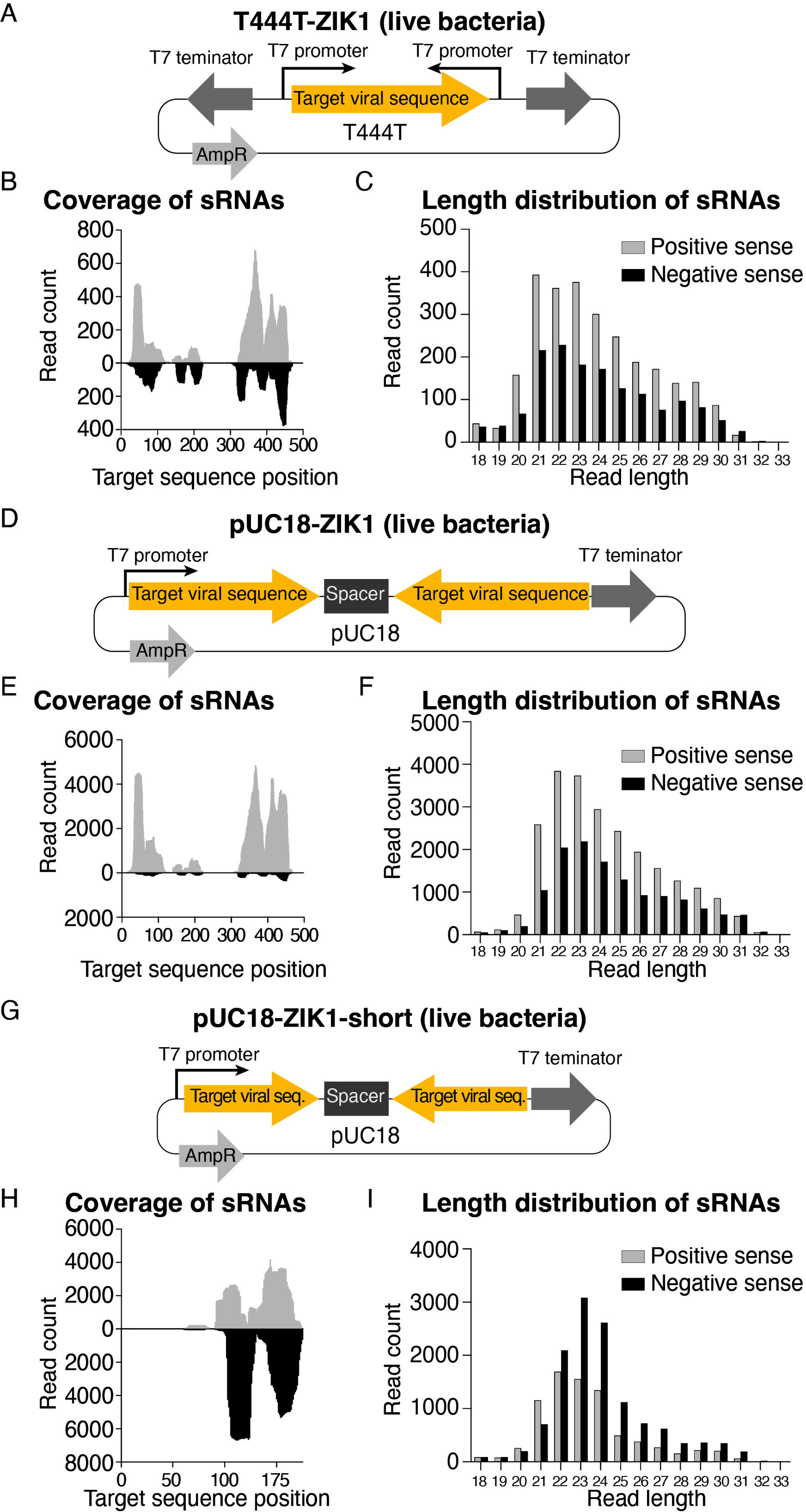
Mosquitoes colonized with dsZIK-producing bacteria do not show production of dsZIK-specific siRNAs. Small RNAs were sequenced from pools of whole mosquitoes colonized for two days with the *E. coli* HT115 strain carrying either the T444T-ZIK1 plasmid (**A**), or the pUC18-ZIK1 plasmid with the 481 nt (**D**) or 175 nt (**G**) ZIKV sequence. For each condition, the number of reads mapping to each position of the positive strand (grey) or negative strand (black) of the target sequence is displayed (**B**, **E**, **H**). The size distribution of the reads shown in (**B, E, H**) is shown in (**C**, **F**, **I**), respectively.

To assess whether different cloning strategies would result in an improved dsRNA-induced response, we colonized mosquitoes with *E. coli* HT115 bacteria carrying the pUC18 hairpin-producing plasmid containing either the 481 bp ZIK1 sequence (**Figure 3D**) or a shorter (175 bp) ZIK1 sequence (**Figure 3G**). We did not find evidence for siRNA production from dsRNA derived from either the long (**Figure 3E-F**) or short construct (**Figure 3H-I**).

To assess whether the observed lack of an antiviral RNAi-based response was due to poor release of dsRNA from live bacteria, we sequenced small RNAs from mosquitoes supplemented with the same *E. coli* strains heat-killed before supplementation, as heat-killed bacteria were previously used to induce oral RNAi in mosquitoes^21^. We did not find evidence for target-specific siRNA production in any of the samples produced from mosquitoes supplemented with heat-killed bacteria (**Supplementary Figure 4A-I**). We conclude that dsRNA produced by bacteria failed to interact with the mosquito RNAi machinery and to induce the production of target-specific siRNAs.

### Oral supplementation of in vitro synthesized dsRNA does not induce an antiviral response in adult mosquitoes

The lack of an interaction between the bacterial-produced dsRNA and the mosquito RNAi machinery could be due to several factors, such as lack of internalization of dsRNA by mosquito gut cells or low stability of dsRNA in the mosquito gut. To investigate whether dsRNA could be internalized by the mosquito gut epithelium and to overcome the potential limiting factor of insufficient release of dsRNA from bacteria, we produced Cy3-labeled dsZIK1 (dsZIK1-Cy3) *in vitro*. Due to the low quantity of dsZIK1-Cy3 that we obtained, we tested whether mosquitoes could feed on a 200 µL sucrose solution provided on a microtube cap (to prevent dilution of dsZIK1-Cy3 in a standard feeding volume). To determine the rate of feeding with this setup, we provided mosquitoes with a sucrose solution containing a blue dye. Mosquitoes that fed on the solution showed a readily visible blue abdomen (**Figure 4A**). We dissected these mosquitoes after one or two days of feeding and verified the presence of blue dye in the crop (where the sugar is stored before digestion) and midgut. While after one day of feeding 67 % (10/15) of mosquitoes showed a blue crop and 80 % (12/15) a blue midgut, after two days all mosquitoes (15/15) were found to have fed on the sucrose solution and showed both a blue crop and midgut. dsZIK1-Cy3 was thus provided to mosquitoes for 48 h in a sucrose solution with the setup described above, but without the blue dye that could potentially interfere with fluorescence detection. Adult mosquitoes fed with dsZIK1-Cy3 were dissected and guts (including crop and hindgut) were observed under a confocal microscope. Mosquitoes fed on sucrose containing dsZIK1-Cy3 showed bright red fluorescence in all gut compartments (**Figure 4B-C**). Interestingly, cells displaying red fluorescence in both the cytoplasm and nucleus were observed in the midgut, which is the region in which blood is stored during digestion and that is primarily infected by viruses ingested with blood (**Figure 4D**). These data indicate that at least some mosquito gut epithelial cells can internalize dsRNA.

**Figure 4.**
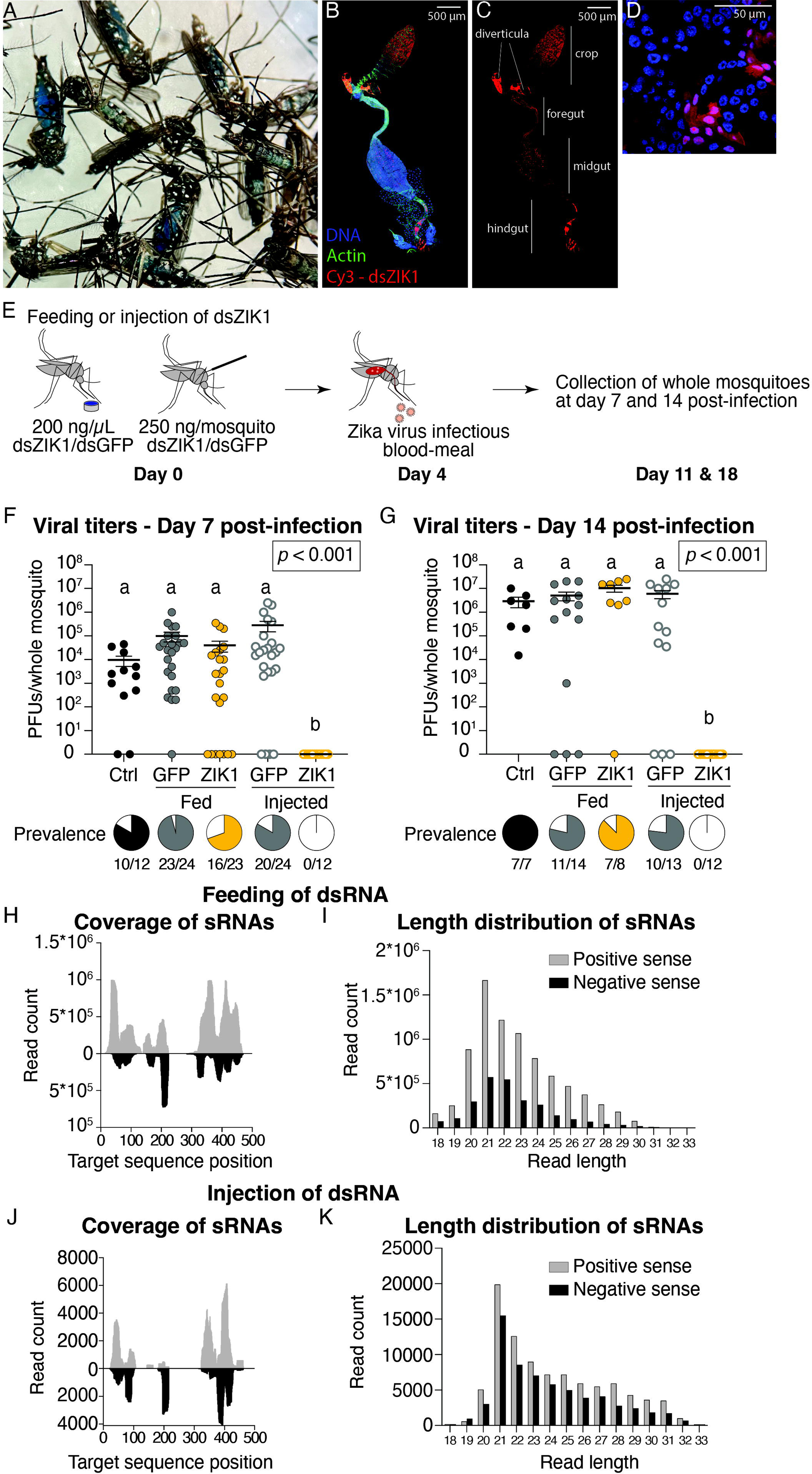
A protective antiviral response is induced by systemic injection of dsRNA in adult mosquitoes, but not by feeding. (**A**) A blue sucrose solution containing dsRNA was provided to mosquitoes for two days. Blue dye in the crop and midgut is visible through the cuticle and allows selection of fed mosquitoes. (**B**-**D**) Confocal images of the digestive tract of a mosquito fed for 48 h with dsZIK1-Cy3. In (**B**) different dyes indicate DNA (blue, DAPI), actin (green, Phalloidin), and dsZIK1-Cy3 (red, Cy3), while in (**C**) only the signal of dsZIK1-Cy3 is shown. In (**C**) the different components of the mosquito digestive tract are indicated. (**D**) represents a higher magnification image of a region of the same midgut shown in (**B**-**C**). Scale bars correspond to 500 µm (**B**-**C**) or 50 µm (**D**). (**E**) Experimental protocol for ZIKV infection of mosquitoes fed or injected with dsZIK1 or dsGFP. At day 0, mosquitoes were fed with a 200 ng/µL solution of dsZIK1 or dsGFP, or injected with 250 ng dsZIK1 or dsGFP/mosquito. After four days, they were offered a ZIKV infectious bloodmeal. At seven and 14 days after infection, whole individual mosquitoes were collected. (**F**-**G**) Viral titers (mean ± SEM) in individual mosquitoes (whole bodies) fed on sucrose (black) or sucrose containing dsGFP (full grey) or dsZIK1 (full yellow), or injected with dsGFP (empty grey) or dsZIK1 (empty yellow). Mosquitoes were collected seven (**F**) or 14 (**G**) days post-infection. Each plot shows the log of plaque forming units (PFUs) measured in individual mosquitoes. Pie charts show the proportion of infected (black/grey/yellow) and uninfected (white) mosquitoes. Numbers below pie charts represent the number of infected over uninfected mosquitoes per tissue/time-point. Viral loads were compared using a Kruskal-Wallis one-way ANOVA followed by a Dunn test with Bonferroni correction. The global ANOVA *p* value is displayed in the top-right corner of each plot and significance groups (*p* < 0.05) are denoted by letters. (**H**-**K**) Small RNAs were sequenced from pools of five whole mosquitoes fed with dsZIK1 for two days (**H**, **I**) or injected with dsZIK1 two days before sampling (**J**, **K**). (**H**, **J**) For each condition, the number of reads mapping to each position of the positive strand (grey) or negative strand (black) of the target sequence is displayed. The size distribution of the reads shown in (**H**, **J**) is shown in (**I**, **K**), respectively.

To explore if *in vitro*-synthesized dsRNA could induce an antiviral RNAi-based response when orally administered to mosquitoes, we fed mosquitoes with non-labeled dsZIK1 and, as a control, with dsGFP (**Figure 4E**). To confirm that the same dsRNA could induce an antiviral effect when introduced systemically into the mosquito body, we injected mosquitoes with dsZIK1 or dsGFP. Four days after the oral supplementation or the injection, we provided mosquitoes with a ZIKV-containing infectious bloodmeal and measured viral titers in whole mosquitoes by plaque assays at seven and 14-days post-infection. We found that oral supplementation of dsZIK1 did not decrease viral titers in mosquitoes collected at either time point, while injection of dsZIK1 completely blocked viral replication at both timepoints (**Figure 4F-G**, Kruskal-Wallis test on log transformed viral titers: day 7 *p* < 0.001; day 14 *p* < 0.001; Kruskal-Wallis test on prevalence: day 7 *p* < 0.001; day 14 *p* < 0.001; see Supplementary Table 2 for individual comparisons). In line with these results, mosquitoes fed with dsCHIK1 did not show any significant reduction in viral titers after infection with CHIKV, while mosquitoes injected with dsCHIK1 showed reduced viral titers in whole bodies two days post-infection and marginally significant reduced titers five days post-infection (**Supplementary Figure 5A-C**, Kruskal-Wallis test on log transformed viral titers: day 2 *p* < 0.001; day 5 *p* = 0.04; Kruskal-Wallis test on prevalence: day 2 *p* = 0.18; day 5 *p* = 0.07; see Supplementary Table 2 for individual comparisons).

To verify that siRNAs were produced from dsRNA introduced into mosquitoes by injection or feeding, we performed small RNA sequencing of RNA from whole mosquito samples. We found that 21 nt siRNAs were enriched in RNA from whole bodies of mosquitoes fed or injected with dsRNA four days before sampling (**Figure 4H-K**).

To investigate the localization and integrity of dsRNA in fed and injected mosquitoes, RNA was extracted from midguts, carcasses, and heads of mosquitoes collected two and four days after feeding/injection (**Supplementary Figure 6A**). Northern blot analysis of samples collected at day two indicated that RNA corresponding to the dsZIK1 sequence was detectable in the midgut, carcass, and head of mosquitoes fed with dsRNA, while only in the carcass and head of mosquitoes injected with dsRNA (**Supplementary Figure 6C**). At day four, a signal from dsZIK1 was detected in the carcass of both fed and injected mosquitoes, and in the midgut of fed mosquitoes (**Supplementary Figure 6C**). It is important to mention that, while the crop was not included in any sample because of the presumable high concentration of dsRNA in this organ, carcass samples contained Malpighian tubules and hindguts, and head samples included the anterior part of the digestive system. Therefore, we cannot determine whether the dsZIK1 signal visible in RNA samples extracted from fed mosquitoes derives from dsRNA that is circulating in the haemocoel and entering other organs or if it represents what is ingested by mosquitoes that fed on dsRNA. However, these results confirmed that dsRNA was not degraded by midgut RNases in fed mosquitoes.

## Discussion

In mosquitoes, targeted RNAi can be activated by introduction of dsRNA corresponding to target sequences. This stimulates the production of target-specific siRNAs from the introduced dsRNA, which are used to guide degradation of complementary RNA. When the introduced dsRNA is designed to target viral RNA, introduction of dsRNA prior to infection can limit subsequent viral replication in a sequence-specific manner. In this paper we tested whether bacteria colonizing the mosquito midgut could be used as a platform to produce virus-specific dsRNA and stimulate an enhanced antiviral response against arboviruses. We showed that the *E. coli* HT115 strain, originally used in *C. elegans* to induce RNAi, is poorly transmitted from mono-colonized larvae to adult mosquitoes but can stably colonize the mosquito gut when provided to adult mosquitoes through sucrose supplementation (**Figure 1D**). However, we did not detect any significant reduction on ZIKV or CHIKV titers in mosquitoes colonized by dsZIK-or dsCHIK-producing bacteria (**Figure 2B-C** and **Supplementary Figure 1E-F**). Moreover, our small RNA sequencing results did not show evidence for the production of 21 nt siRNAs specific for the viral target sequence in mosquitoes provided with live or heat-killed dsRNA-producing bacteria (**Figure 3**, **Supplementary Figure 4**). We identified at least three potential issues that may have limited the interaction of the bacterial-produced dsRNA with the mosquito RNAi machinery: (i) insufficient production or release of dsRNA by bacteria in the mosquito gut, (ii) low stability of dsRNA in the mosquito gut, and/or (iii) limited uptake of dsRNA from mosquito gut cells.

Although we confirmed dsRNA production *in vivo* by bacteria (**Supplementary Figure 2**), we cannot confirm whether dsRNA was released by bacteria in the mosquito gut. In previous experiments in *C. elegans*, *E. coli* HT115 was used as a source of food for the nematode and undergoes cell lysis during ingestion and release dsRNA^22,33^. It is unknown to what extent bacterial cells are lysed in the mosquito gut. Live bacteria are needed at the larval stage to support mosquito development and are found in the larval gut by two hours post-hatching, but dead bacteria can be found in the larval gastric caeca and midgut by 16 hours post-hatching^34^. This suggests that, at least at the larval stage, some bacteria undergo cell lysis in the mosquito gut and potentially release dsRNA. Live, culturable bacteria have been isolated from the adult gut from both lab-reared and field-collected mosquitoes^35–37^ and an expansion of some members of the microbiota is observed 24 h after a bloodmeal^38,39^. However, the relative proportions of live and dead bacteria in adult mosquitoes are unknown.

To test whether production or release of dsRNA from bacteria was a limiting factor in our system, we decided to orally administer naked dsRNA to mosquitoes. Northern blot analysis of different mosquito tissues confirmed that dsRNA molecules were, at least in part, intact after being provided to mosquitoes via oral supplementation for two days, indicating that orally acquired dsRNA is not totally degraded by gut RNases (**Supplementary Figure 6C**). Although oral acquisition of dsZIK induced production of dsZIK-derived siRNAs in mosquitoes (**Figure 4H-I**), it failed to trigger an antiviral RNAi response and limit ZIKV replication (**Figure 4F-G**). In contrast, the same dsZIK provided systemically though injection completely blocked viral replication (**Figure 4F-G**). This suggests that dsRNA provided via oral supplementation is not able to induce an RNAi response capable of inhibiting viral replication. Downstream from siRNA production, RNAi has been shown to have a limited role in restricting DENV or ZIKV infection in the *Ae. aegypti* midgut, predominantly due to the lack of expression of the dsRNA-binding protein, Loqs2, in this tissue^40^. This may explain why siRNA produced after oral administration of dsRNA failed to produce an effective RNAi-based response. To overcome this limitation, further applications involving oral RNAi should explore the co-expression of Loqs2 in the mosquito midgut.

Cy3-labeled dsRNA provided orally to mosquitoes could be visualized in the crop, diverticula, and some midgut cells, where fluorescence could be detected in both the cytoplasm and nucleus (**Figure 4B-D**). However, we did not investigate whether orally supplemented dsRNA could be found in other tissues distal from the digestive system. A recent study described a similar strong fluorescence signal in the alimentary canal of *Ae. aegypti* larvae and adults fed with fluorescently-labeled dsRNA^41^. Interestingly, this study showed that adult mosquitoes fed with fluorescently-labeled dsRNA did not display fluorescence in other tissues, while mosquitoes intrathoracically injected with dsRNA showed strong fluorescence in hemocytes, pericardial cells, and ovaries. Uptake of dsRNA and amplification of the RNAi signal by hemocytes is essential for effective antiviral RNAi in *Drosophila melanogaster*^42^. We can therefore speculate that the failure of orally acquired dsRNA to inhibit viral replication in *Ae. aegypti* may be due to incorrect processing of this dsRNA, which may be internalized by midgut cells, but not hemocytes or cells in other tissues relevant for an antiviral response.

The only previous report of successful microbiota-mediated antiviral RNAi in insects was in honeybees infected with deformed wing virus, where bees colonized by viral dsRNA-producing microbiota were found to exhibit longer lifespans and reduced viral replication compared to bees carrying dsGFP-producing bacteria^8^. However, mosquitoes and bees might have physiological differences that could limit this application against arboviruses. For instance, the honeybee microbiota almost exclusively colonizes the hindgut^43^, while mosquitoes have high bacterial loads mainly in the crop and midgut^44^. Differences in viral infection dynamics and/or dsRNA uptake by different tissues might explain the different results obtained with *Apis millifera* and *Ae. aegypti*.

Over the last ten years, oral RNAi targeting essential mosquito genes has been proposed as a vector control strategy to inhibit or block mosquito development and/or reproduction. Several methods have been developed to deliver dsRNA to mosquito larvae or adults, including feeding of naked dsRNA, colonization with microorganisms producing dsRNA, or feeding with nanoparticle-dsRNA complexes (reviewed in ^21^). However, some studies have reported high variability or low gene silencing efficiency when dsRNA was provided via oral feeding^15,16^ and it is likely that other studies with contradictory or negative results were left unpublished. As emphasized in a recent discussion by Devang Mehta, the importance of showcasing negative results for the advancement of science cannot be overstated^45^. It is key to acknowledge that when negative results go unpublished, other scientists can’t learn from them and end up repeating failed experiments, leading to a waste of public funds and a delay in genuine progress. Our results point to clear limitations of oral RNAi as a strategy to disrupt viral transmission by mosquitoes and should be considered when developing new strategies for vector control.

## Materials and Methods

### Bacteria, viruses, cells and mosquitoes

*E. coli* HT115(DE3, Genotype: F-, mcrA, mcrB, IN(rrnD-rrnE)1, rnc14::Tn10) was obtained from the Caenorhabditis Genetics Center (CGC, University of Minnesota). For all experiments, bacteria were inoculated from fresh colonies in Luria-Bertani (LB) broth supplemented with 20 µg/mL tetracycline. The growth medium was supplemented with 100 µg/mL ampicillin for bacteria carrying the T444T or pUC18 plasmids.

The ZIKV strain used in this study is the prototype African MR766 strain, derived from a previously described infectious clone^46^. The CHIKV strain used is the Caribbean strain, Asian genotype originally described in^47^. Viral stocks were produced in Vero E6 cells.

Vero E6 cells were cultured in Dulbecco’s modified Eagle’s medium (DMEM, Life Technologies) supplemented with 10 % fetal bovine serum (Invitrogen) and 1 % Penicillin/Streptomycin (P/S, Life Technologies) at 37 °C with 5 % CO_2_.

*Ae. aegypti* mosquitoes belonged to the 28^th^–31^st^ generation of a colony created in 2013 from mosquitoes collected in the Kamphaeng Phet Province, Thailand^48^. They were maintained under standard insectary conditions (27 °C, 70 % relative humidity, 12h:12h light/dark cycle). Mosquitoes are routinely blood-fed with rabbit blood with an Hemoteck system (Hemotek Ltd). Conventionally reared larvae were reared in 30×40 cm plastic trays with 1.5 L of dechlorinated tap water and ground TetraMin fish food. Pupae were transferred in plastic cages (BugDorm) for emergence and adults were provided with autoclaved 10 % sucrose solution.

### Selection of viral target sequences for dsRNA production

Sequences of the ZIKV (MR766 strain) or CHIKV (Caribbean strain) genome spanning regions of the polyprotein coding sequence corresponding to different mature proteins were selected and used as a blastn^49^ query against the *Ae. aegypti* genome in Vectorbase and all *Aedes* sequences in Genebank to verify that they did not target any *Aedes* sequence. To exclude any interference by mosquito small RNAs on the target viral sequence, small RNA library data previously produced in the lab from *Ae. aegypti* uninfected mosquitoes were mapped on the ZIKV and CHIKV sequences. Any mosquito small RNA was found to map on the selected viral sequences.

### Cloning

The T444T plasmid was a gift from Tibor Vellai (Addgene plasmid #113081; RRID: Addgene_113081^23^). GFP and target viral sequences were cloned into T444T using *Not*I and *Age*I restriction sites and standard cloning techniques. The NEBuilder® HiFi DNA Assembly Kit (New English Biolabs) was used to insert the following elements into the pUC18 plasmid: the T7 promoter, *Not*I and *Age*I restriction sites, a spacer sequence corresponding to a 300 bp portion of an intron of the *Drosophila melanogaster actin* gene (GenBank: M18829.1), *Nhe*I and *Kpn*I restriction sites and the T7 terminator sequence. The final pUC18-spacer plasmid sequence and map are available upon request. The same primers used for the cloning in T444T were used to clone GFP and target viral sequences in pUC18-spacer using *Not*I and *Age*I restriction sites. To clone the inverted repeat, *Nhe*I and *Kpn*I restriction sites were used. A shorter ZIK1 sequence (175 bp) was also cloned in pUC18-spacer using the same cloning strategy. All primers used are listed in Supplementary Table 1. After cloning, plasmid sequences were confirmed via Sanger sequencing from both sides of the insert.

### Mono-colonization of Ae. aegypti larvae with E. coli HT115

Egg sterilization and larval mono-colonization were performed in sterile conditions working inside a microbiological safety cabinet. *Ae. aegypti* eggs were surface sterilized with subsequent 5-minute washes in 70 % ethanol, 1 % bleach, and 70 % ethanol. After three rinses in sterile Milli-Q water, eggs were transferred to a 25 mL cell culture flask with vented cap overnight for hatching. On the same day, *E. coli* HT115 carrying the selected plasmid was used to inoculate LB supplemented with 40 µM IPTG, 20 µg/mL tetracycline and 100 µg/mL ampicillin, and grown for ∼ 16 h at 37 °C, 200 rpm shaking. The following day, the bacterial culture was centrifuged and resuspended in sterile Milli-Q water at a final concentration of 10^8^ CFUs/mL. First instar larvae were placed in plastic boxes cleaned with ethanol (200 larvae per box), together with 500 mL of bacterial suspension, 40 µM IPTG and 5 mL of sterile fish food (TetraMin baby, 50 mg/mL). Boxes were closed with filtered covers and kept at 27 °C, 12h:12h light/dark cycle. Pupae were transferred into sterile boxes for emergence as in^32^. Larval water was plated on LB agar and LB agar supplemented with 100 µg/mL ampicillin to confirm the mono-colonization of mosquitoes. The same number of colonies and the same colony morphology were observed in both media.

To monitor the transstadial transmission of *E. coli* HT115-T444T-ZIK1 from larvae to adults, adult mosquitoes originating from mono-colonized larvae were either provided with 10 % sterile sucrose or with bacteria resuspended in 10 % sterile sucrose at a final concentration of 10^9^ CFUs/mL. At days one and four post-emergence, mosquitoes were collected, surface sterilized for 3 minutes in 70 % ethanol, rinsed in sterile PBS, and individually homogenized in 200 µL LB. Mosquito homogenates were serially diluted in LB and plated on LB agar and LB agar supplemented with 100 µg/mL ampicillin. Plates were placed overnight at 37 °C and the number of colonies was counted to estimate whole mosquito bacterial loads.

### Infection of mono-colonized mosquitoes with ZIKV or CHIKV

Larvae mono-colonized by the selected strain of *E. coli* HT115 were produced as described above. Larval water was supplemented with 40 µM IPTG to induce the transcription of the dsRNA by bacteria. Adult mosquitoes were provided with a 10 % sterile sucrose solution supplemented with 10^9^ CFUs/mL bacteria and 40 µM IPTG for four days. The day before the infectious bloodmeal, mosquitoes were deprived of the sucrose/bacteria/IPTG solution for starvation. Mosquitoes were transferred to a biosafety level 3 insectary and offered an infectious bloodmeal on a Hemoteck system (Hemotek Ltd) consisting of washed rabbit blood supplemented with 10^6^ PFUs/mL of ZIKV or CHIKV and 10 mM ATP. Fully engorged mosquitoes were kept at 27 °C, 70 % relative humidity, 12h:12h light/dark cycle until sampling and provided with fresh bacteria and sucrose every three days. At the selected time points, mosquitoes were dissected and tissues were placed in 200 µL PBS. Samples were kept at −80 °C until processing.

### Plaque assay

The day before the assay, Vero-E6 cells were seeded in 24-well plates at a density of 1.75*10^5^ cells/well in DMEM with 10 % FBS and 1 % P/S. The day of the assay, mosquito samples were homogenized in a bead beater for 30 sec at 5000 rpm. Ten-fold dilutions of mosquito homogenates or of an aliquot of the bloodmeal were prepared in DMEM. Two-hundred µL of each dilution were added to the cell monolayer and incubated for 1 h at 37 °C with 5 % CO_2_. Cells were covered with DMEM with 2 % FBS, 1 % P/S and 0.8 % agarose. After seven (ZIKV) or three (CHIKV) days, cells were fixed with 4 % paraformaldehyde. Plaques were counted after 0.2 % crystal violet staining.

### Small RNA sequencing

RNA was extracted from pools of five mosquitoes using TRIzol. Small RNAs of 18-33 nt in length were purified from a 15% acrylamide/bisacrylamide (37.5:1), 7 M urea gel as described previously^50^. Libraries were prepared using the NEBNext® Small RNA Library Prep Set for Illumina® (New England BioLabs), with the Universal miRNA Cloning Linker (New England BioLabs) as the 3’ adaptor and in-house designed indexed primers. Libraries were diluted to 4 nM and sequenced using an Illumina NextSeq 500 High Output kit v2 (75 cycles) on an Illumina NextSeq 500 platform.

### Bioinformatics analysis of small RNA libraries

FastQC^51^ was used to assess the quality of fastq file. Low quality portions and adaptors were trimmed from each read using cutadapt^52^. Only reads with a phred score ≥20 were kept. FastQC was used on the fastq files created by cutadapt to evaluate the overall quality. Mapping was performed using Bowtie^53^ with the -v 1 option (one mismatch between the read and its target) using as reference sequences the ZIKV, CHIKV, or GFP target sequences, or the PCLV genome. Bowtie generates results in sam format. All sam files were analyzed by different tools of the package samtools^54^ to produce bam indexed files. To analyze these bam files, different kind of graphs were generated using home-made R scripts with several Bioconductor libraries such as Rsamtools and Shortreads^55^.

### Synthesis and labelling of dsRNA

RNA corresponding to the same ZIK1, CHIK1 and GFP sequences cloned in the T444T plasmid was transcribed *in vitro* using the T7 MEGAscript Kit (Invitrogen) following the manufacturer’s instructions. A PCR amplicon generated with both forward and reverse primers containing the T7 promoter sequence (Supplementary Table 1) was used as a template for the *in vitro* transcription. A DNase treatment was performed after transcription to eliminate the DNA template. The annealing of the two strands of dsRNA was allowed by heating the RNA at 98 °C for 5 min and leaving it to cool down slowly to room temperature. After quantification with Nanodrop, RNA was stored at −80 °C until use. The Silencer™ siRNA Labeling Kit with Cy™3 dye (Invitrogen) was used to label dsRNA following the manufacturer’s instructions. An aliquot of the labeled dsRNA was run on a 1 % agarose gel to visualize a shift in the migration and thus confirm the labeling.

### Oral feeding of mosquitoes with labeled dsRNA, sample collection and visualization

One-to-three day old conventionally reared female mosquitoes were starved for one day and then provided with a 10 % sucrose solution containing 200 ng/µL of the labeled dsRNA. After two days of feeding, midguts were dissected and fixed overnight in 4 % methanol-free paraformaldehyde in PBS. After three washes with PBS, samples were permeabilized in 1 % BSA, 0.1 % Triton X-100 in PBS for 1 h at room temperature. After three washes with PBS, midguts were incubated for 30 minutes with DAPI and Oregon Green 488 Phalloidin (Invitrogen) dyes to stain DNA and actin, respectively. After three additional washes in PBS, midguts were mounted on slides with Vectashield H-1000 (Vector Laboratories) and visualized on a Zeiss LSM 780 confocal microscope at 100X magnification. To reconstruct mosquito full digestive tract images, multiple flanking images were acquired using the “tile” function and subsequently assembled using the ZEN Microscopy Software.

### Infection of mosquitoes fed or injected with dsRNA

One-to-three day old conventionally reared female mosquitoes were starved for one day and then provided with a 10 % sucrose solution containing 200 ng/µL of dsRNA and 1 mg/mL of the blue dye, Erioglaucine disodium salt (Sigma-Aldrich). The control consisted of mosquitoes fed on blue-dyed sucrose without dsRNA. All mosquitoes were allowed to feed on the sucrose solution for three days and then only mosquitoes showing a blue abdomen were selected for infection. In parallel, one-to-three day old conventionally reared female mosquitoes were intrathoracically injected with 250 ng of dsRNA using a Nanoject III system (Drummond). They were left to recover for three days. The day before the infectious bloodmeal, mosquitoes were deprived of sucrose for starvation. Mosquitoes were infected with 10^6^ PFUs/mL of ZIKV MR766 or CHIKV as described above. At the selected time points, whole mosquitoes were collected in 2 mL screwcap tubes containing 200 µL of sterile PBS and glass beads. Samples were kept at −80 °C until processing. Viral titers in whole mosquitoes were measured by plaque assay as described above.

### Northern blot

ZIK1 sense and antisense RNA probes labeled with digoxigenin were synthesized using the DIG Northern Starter Kit (Roche) following the manufacturer’s instructions. The template consisted of a PCR product obtained with primers containing the T7 (sense) or T3 (antisense) promoter sequence (Supplementary Table 1). Serial dilutions of the two probes were visualized on a dot blot for probe quantification.

*E. coli* HT115-T444T-ZIK1 or-CHIK1 (probe specificity control) was inoculated in 5 mL LB supplemented with 20 µg/mL tetracycline and 100 µg/mL ampicillin and grown overnight at 37 °C shaking at 200 rpm. The following day, 100 µL of the overnight culture were used to inoculate 5 mL of LB with 20 µg/mL tetracycline and 100 µg/mL ampicillin with or without 40 µM IPTG. Bacteria were grown overnight at 37 °C shaking at 200 rpm. After 24 h, 1 mL of bacterial culture was centrifuged and the bacterial pellet was resuspended in 100 µL of 0.1 % SDS. Bacterial extracts were incubated for 2 min at 95 °C for cell lysis and RNA was extracted using TRIzol.

Mosquitoes fed or injected with dsRNA were dissected after two or four days of feeding with 200 ng/µL dsZIK or after injection with 250 ng/mosquito of dsZIK1. For each time-point/condition, 15 to 28 mosquitoes were dissected and their midguts, carcasses (excluding crop, including hindgut and Malpighian tubules), and heads were pooled and stored at −80 °C until processing. Total RNA was extracted from each sample pool using TRIzol.

For northern blot analysis, RNA samples were mixed with a glyoxal-containing loading dye, incubated at 50 °C for 30 min for RNA denaturation, and loaded on a 1.2 % agarose gel. Five µg of total RNA were loaded on the gel for all samples (including *in vitro* synthesized dsRNA) except for midguts at day 4 (injected) and heads (fed/injected) at day 4, for which 1.5 µg of RNA were loaded due to low RNA yields. After gel electrophoresis, RNA was transferred to a Nytran Super Charge membrane (Whatman) using the NorthernMax^TM^-Gly kit (Invitrogen) following the manufacturer’s instructions. After transfer and crosslinking with UV light, the membrane was incubated overnight with 0.2 ng/µL of the selected probe. Membranes were incubated with an Alkaline Phosphatase-conjugated Anti-Digoxigenin antibody and the chemiluminescent signal was revealed using a ChemiDoc MP imaging system (Biorad).

### Statistics

All graphs were created using GraphPad Prism (version 9.5.1) or R (version 4.2.1). Statistical analyses were performed in R (version 4.2.1). The normality of data was tested with a Shapiro test. A Wilcoxon Signed Rank test was used to compare mosquito bacterial loads, prevalence and log_10_ transformed viral titers of mosquitoes colonized by dsZIK-producing bacteria. A Kruskal-Wallis one-way analysis of variance followed by a Dunn test with Bonferroni correction was used to compare log_10_ transformed viral titers in mosquitoes colonized by dsCHIK producing bacteria, or in mosquitoes fed or injected with dsZIK/dsCHIK. Detailed results of statistical analyses are shown in Supplementary Table 2. R codes are available upon request.

## Data availability

The small RNA sequencing data were deposited to the Sequence Read Archive under accession number PRJNA1034089.

## Acknowledgments

We are grateful to Jared Nigg and Louis Lambrechts for critically reading of the manuscript. We thank Catherine Lallemand for technical assistance in mosquito rearing, and Madeleine Lausted for assistance in dsRNA feeding.

This work was supported by funding from the French Government’s Investissement d’Avenir program, Laboratoire d’Excellence Integrative Biology of Emerging Infectious Diseases (grant ANR-10-LABX-62-IBEID), Agence Nationale de la Recherche (grant ANR-22-CE35-0001, MAMMAMIA), and Fondation iXcore - iXlife - iXblue Pour La Recherche to M.C.S., the Pasteur-Roux-Cantarini fellowship of Institut Pasteur to O.R. We acknowledge the kind financial support to the Photonic BioImaging (UTechS PBI) platform by the Institut Pasteur (Paris), the France–BioImaging infrastructure network supported by the Agence Nationale de la Recherche (ANR-10–INBS–04, Investments for the future), and the Région Ile-de-France (program DIM-Malinf).

## Authors contributions

O.R. and M.-C.S. designed the experiments; O.R., A.H.-L., and H.B. performed the experiments; O.R. and L.F. analyzed the data; O.R. wrote the paper with input from co-authors. All authors reviewed and approved the final version of the manuscript.

## Supplementary figures and tables legends

**Supplementary Figure 1. Mosquitoes colonized with dsCHIK-producing bacteria do not show reduced viral titers.** (**A**) Representation of the T444T plasmid. The target viral sequence was cloned between two directionally opposed T7 promoters to allow convergent transcription of a sense and antisense RNAs that form dsRNA upon annealing. (**B**) Representation of the pUC18 plasmid. An inverted repeat of the target viral sequence was cloned downstream from a T7 promoter, allowing the formation of an RNA hairpin after transcription. (**C**) Position of the target viral sequence on the CHIKV genome. The selected sequence is 557 nt long and spans the coding sequences for the Capsid (Cap) and the Envelope (E3) peptide coding genes. The other components of the CHIKV genome are NSPs (Non-Structural Proteins). (**D**) Experimental protocol for CHIKV infection of mosquitoes colonized by dsRNA-producing *E. coli*. Mosquitoes originating from mono-colonized larvae were kept on bacteria-supplemented sucrose for four days. After one day of starvation, they were offered a CHIKV-containing infectious bloodmeal and subsequently kept with bacteria-supplemented sucrose until sampling at two and five days post-infection. At each time point, bodies and heads of individual mosquitoes were collected. (**E-F**) Viral titers (mean ± SEM) in bodies and heads of individual mosquitoes colonized by *E. coli* HT115 carrying the T444T-GFP (grey), T444T-CHIK1 (dark blue), or pUC18-CHIK1 (light blue) plasmid and collected two (**E**) or five (**F**) days post-infection. Each plot shows the log of plaque forming units (PFUs) measured in individual mosquitoes in each tissue. Pie charts show the proportion of infected (grey/blue) and uninfected (white) mosquitoes. Numbers below pie charts represent the number of infected over uninfected mosquitoes per tissue/time-point. Viral loads were compared using a Kruskal-Wallis one-way ANOVA followed by a Dunn test with Bonferroni correction. The global ANOVA *p* value is displayed in the top-right corner of each plot and significance groups (*p* < 0.05) are denoted by letters.

**Supplementary Figure 2. Northern blot analysis of *E. coli* HT115-T444T-ZIK1 total RNA.** (**A**, **B**) Ethidium bromide-stained agarose gel (**A**) and northern blot (**B**) of dsZIK1 synthesized *in vitro* and total RNA extracted from a 24 h bacterial culture without and with 40 µM IPTG induction. For (**B**), the same samples were loaded in duplicate on the same agarose gel and transferred on the same membrane. The membrane was cut and incubated with either a sense (left) or antisense (right) 345 nt probe annealing to the ZIK1 sequence. (**C**, **D**) Specificity of the ZIK1 sense probe. Ethidium bromide-stained agarose gel (**C**) and northern blot (**D**) of the same samples showed in (**A**, **B**) loaded together with dsCHIK1 synthesized *in vitro* and total RNA extracted from a 24 h culture of *E. coli* HT115-T444T-CHIK1 without and with 40 µM IPTG induction. Northern blot membrane was incubated with the sense ZIK1 probe to show binding specificity to dsZIK1 RNA.

**Supplementary Figure 3. Sequencing quality control of small RNAs mapping on the Phasi-Charoen Like Virus (PCLV) genome.** Small RNAs were sequenced from the pools of whole mosquitoes shown in Figure 3 and mapped to the PCLV genomic segments S (**A**-**B**), M (**C**-**D**), and L (**E**-**F**). The number of reads mapping to each position of the positive strand (grey) or negative strand (black) of the PCLV genome is shown (**A**, **C**, **E**). The size distribution of the reads shown in (**A, C, E**) is shown in (**B**, **D**, **F**), respectively.

**Supplementary Figure 4. Mosquitoes colonized with heat-killed dsZIK-producing bacteria do not show production of dsZIK-specific siRNAs.** Small RNAs were sequenced from pools of whole mosquitoes supplemented for two days with sucrose containing heat-killed *E. coli* HT115 (heated for 5 min at 98 °C) carrying either the T444T-ZIK1 plasmid (**A**), or the pUC18-ZIK1 plasmid with the 481 nt (**D**) or 175 nt (**G**) ZIKV sequence. For each condition, the number of reads mapping to each position of the positive strand (grey) or negative strand (black) of the target sequence is displayed (**B**, **E**, **H**). The size distribution of the reads shown in (**B, E, H**) is shown in (**C**, **F**, **I**), respectively.

**Supplementary Figure 5. Effect of oral supplementation or injection of *in vitro* synthesized dsRNA on CHIKV infection.** (**A**) Experimental protocol for CHIKV infection of mosquitoes fed or injected with dsCHIK1 or dsGFP. At day 0, mosquitoes were fed with a 200 ng/µL solution of dsCHIK1 or dsGFP, or injected with 250 ng dsCHIK1 or dsGFP/mosquito. After four days, they were offered a CHIKV infectious bloodmeal. At two and five days after infection, whole individual mosquitoes were collected. (**B-C**) Viral titers (mean ± SEM) in individual mosquitoes (whole bodies) fed on sucrose (black) or sucrose containing dsGFP (full grey) or dsCHIK1 (full blue), or injected with dsGFP (empty grey) or dsCHIK1 (empty blue). Mosquitoes were collected two (**B**) or five (**C**) days post-infection. Each plot shows the log of plaque forming units (PFUs) measured in individual mosquitoes. Pie charts show the proportion of infected (black/grey/blue) and uninfected (white) mosquitoes. Numbers below pie charts represent the number of infected over uninfected mosquitoes per tissue/time-point. Viral loads were compared using a Kruskal-Wallis one-way ANOVA followed by a Dunn test with Bonferroni correction. The global ANOVA *p* value is displayed in the top-right corner of each plot and significance groups (*p* < 0.05) are denoted by letters.

**Supplementary Figure 6. Northern blot analysis of total RNA extracted from mosquitoes fed or injected with dsZIK1.** (**A**) Experimental protocol for sample collection of mosquitoes fed or injected with dsZIK1. At day 0, mosquitoes were fed with a 200 ng/µL solution of dsZIK1 or injected with 250 ng dsZIK1/mosquito. After two and four days, midguts, carcasses, and heads of individual mosquitoes were collected and pooled. RNA was extracted from pools of 15 to 28 mosquitoes. (**B**-**C**) Ethidium bromide-stained agarose gel (**B**) and northern blot (**C**) of RNA samples extracted from pools of midguts (M), carcasses (C), or heads (H) of mosquitoes fed or injected with dsZIK1 collected two or four days after feeding/injection. The membrane was incubated with a sense 345 nt probe annealing to the ZIK1 sequence. dsZIK1 synthesized *in vitro* was loaded on the same gel as a control.

**Supplementary Table 1. List of primers used in this study.** For each primer, the following information is indicated: name, sequence (5’-3’), GenBank reference sequence, position on the reference sequence, amplicon length and application.

**Supplementary Table 2. Results of statistical analyses.**

