## Supplementary figures and images for "Challenges in the Biotechnological Implementation of Oral RNA Interference as an Antiviral Strategy in *Aedes aegypti*"

### Supplementary Figure 1

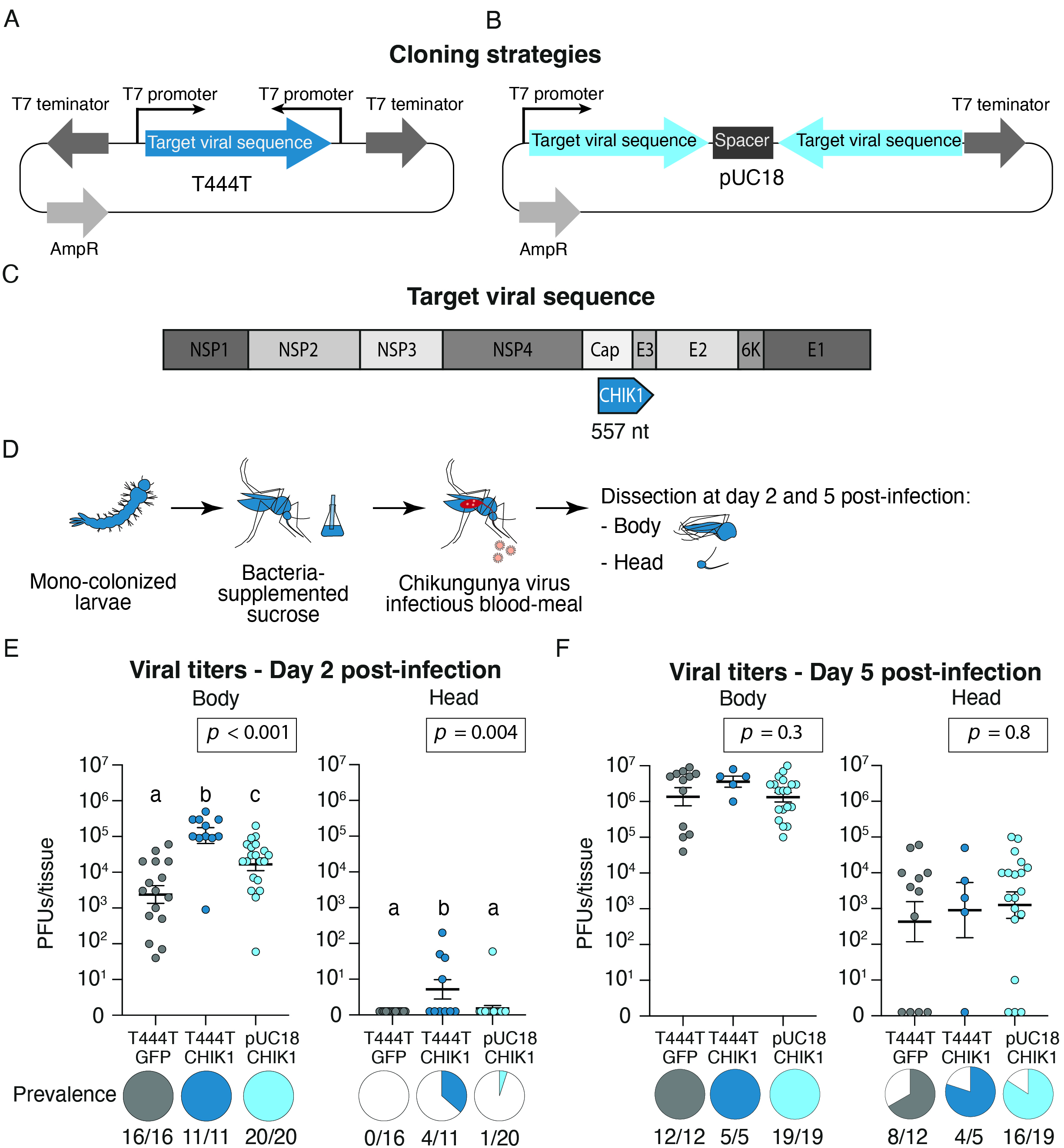

### Supplementary Figure 2

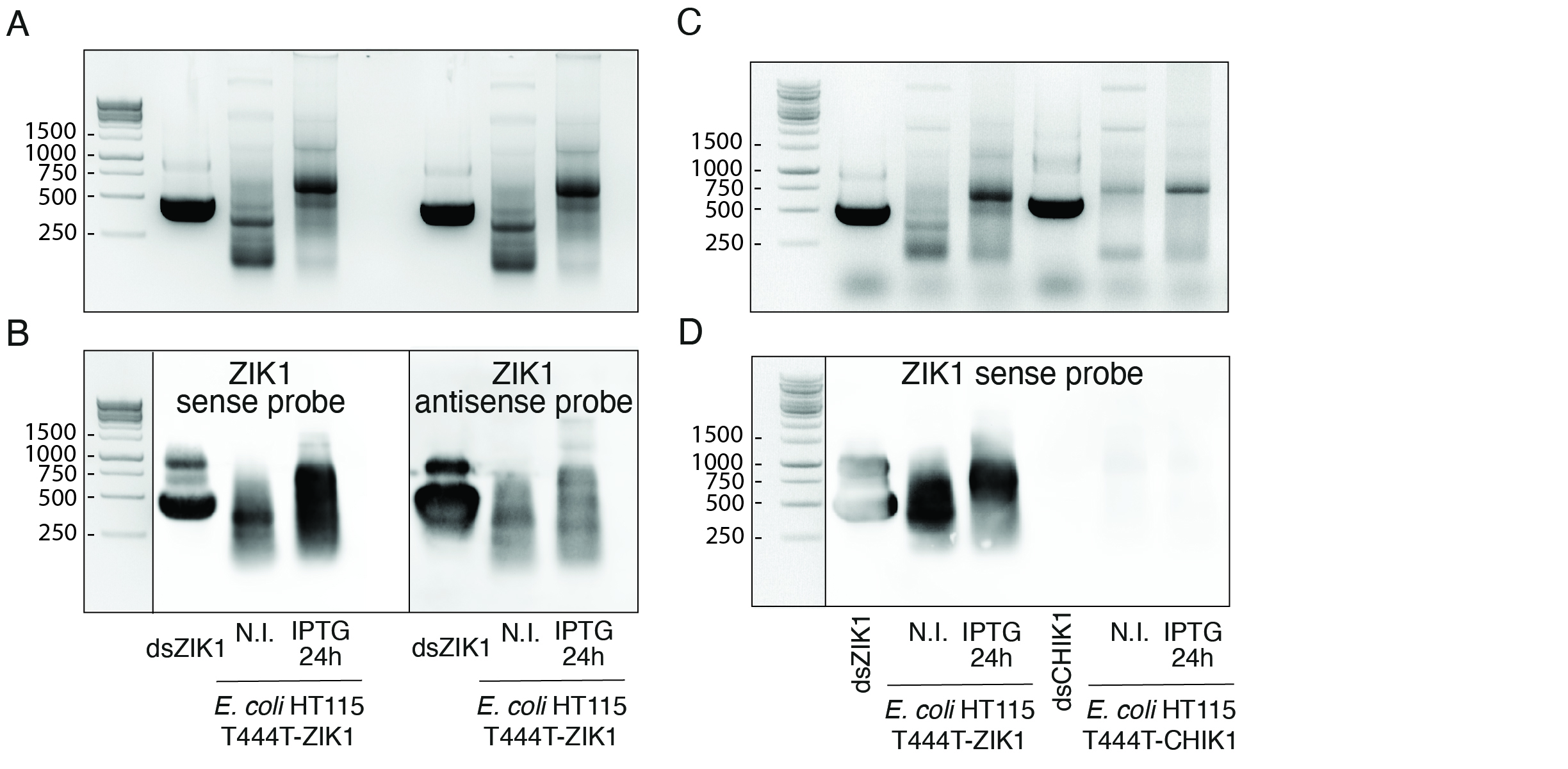

### Supplementary Figure 3

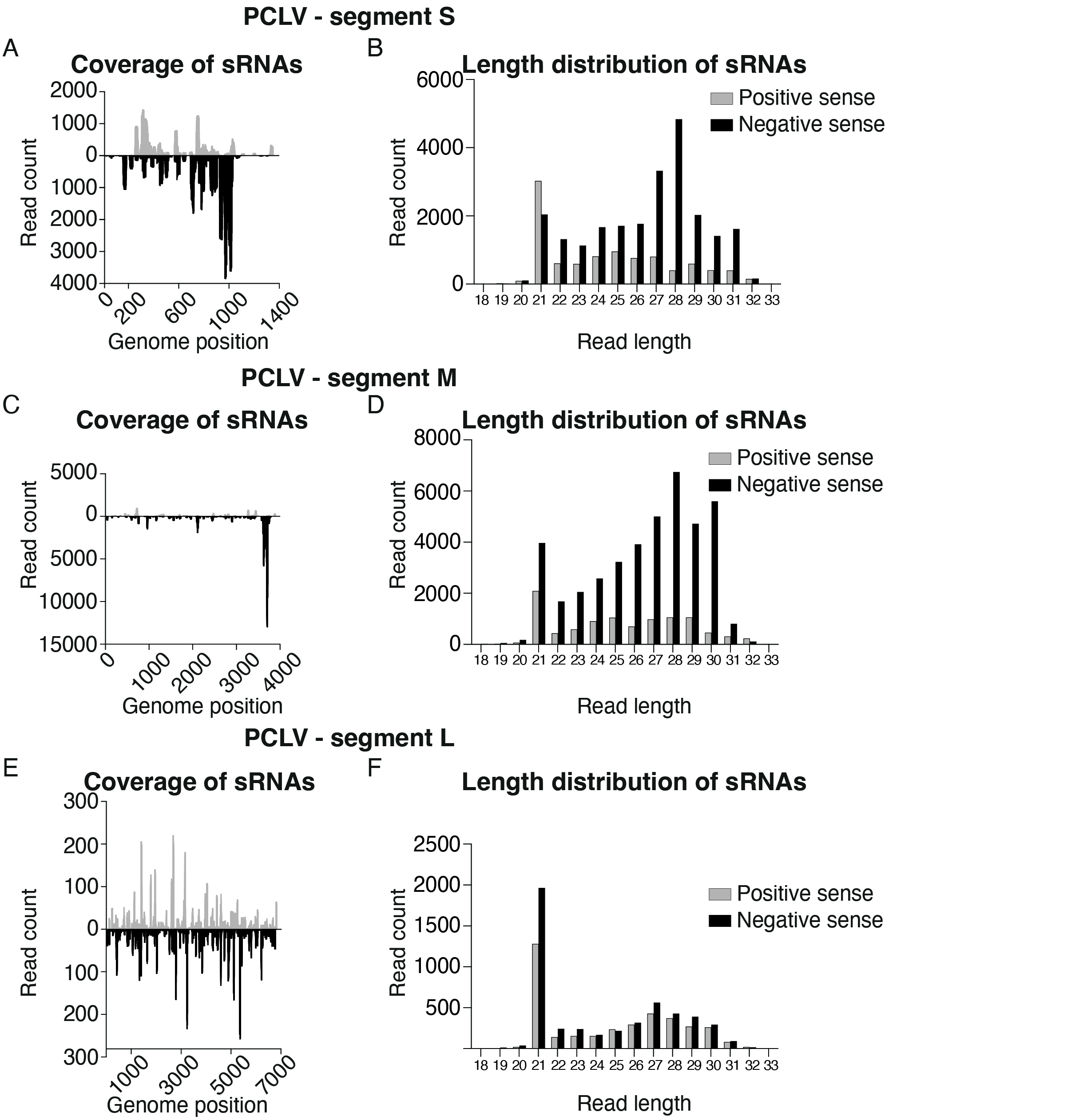

### Supplementary Figure 4

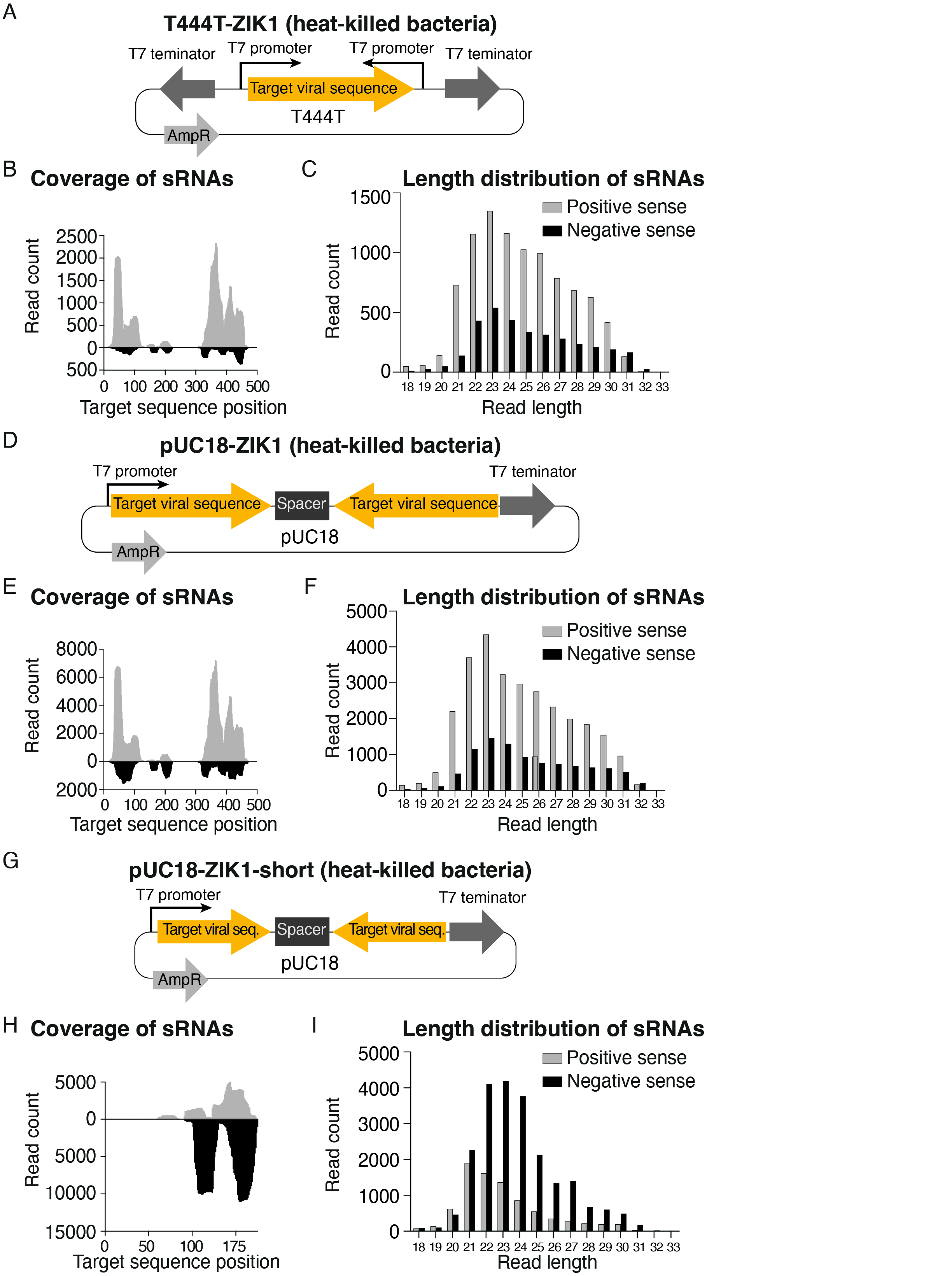

### Supplementary Figure 5

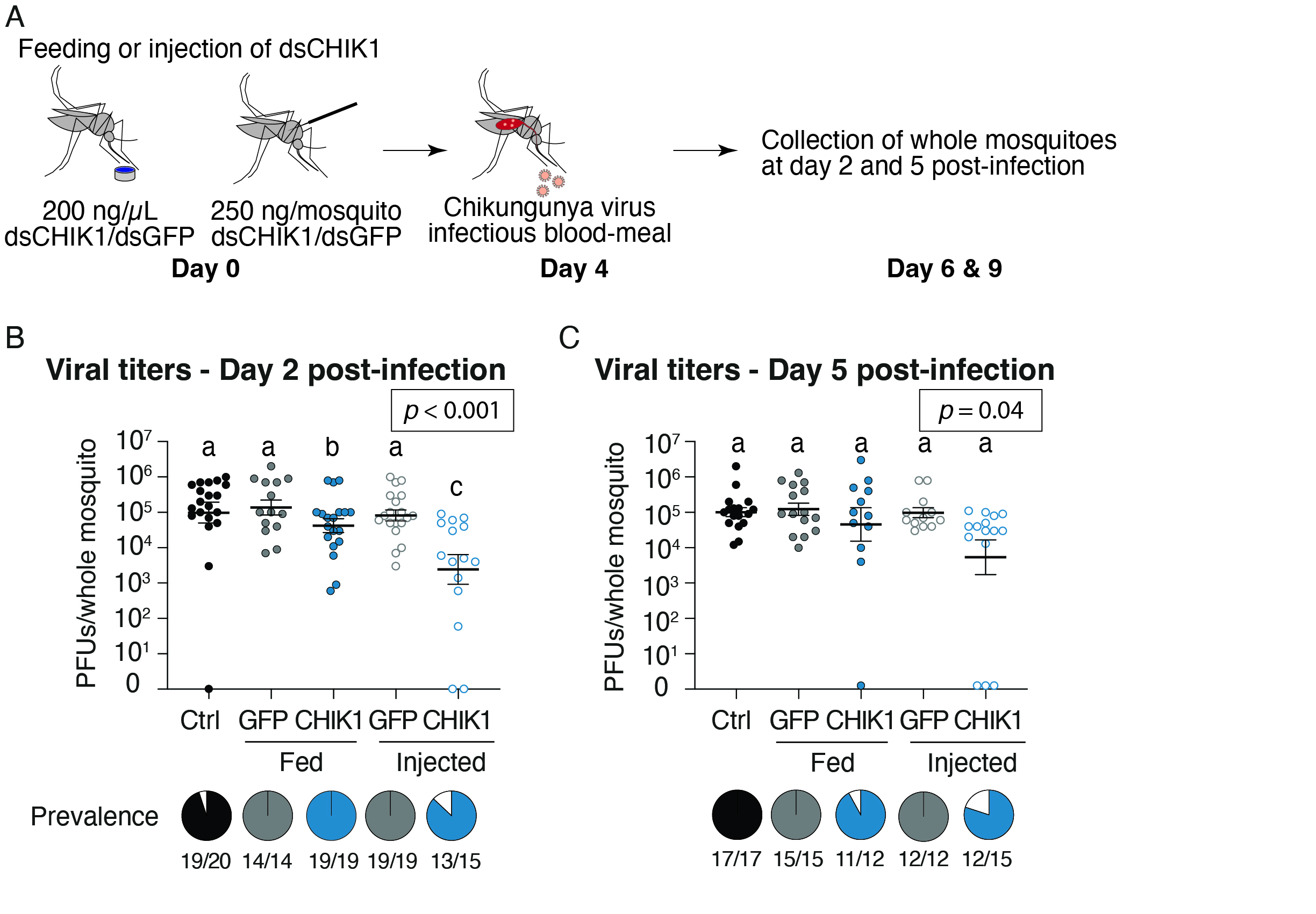

### Supplementary Figure 6

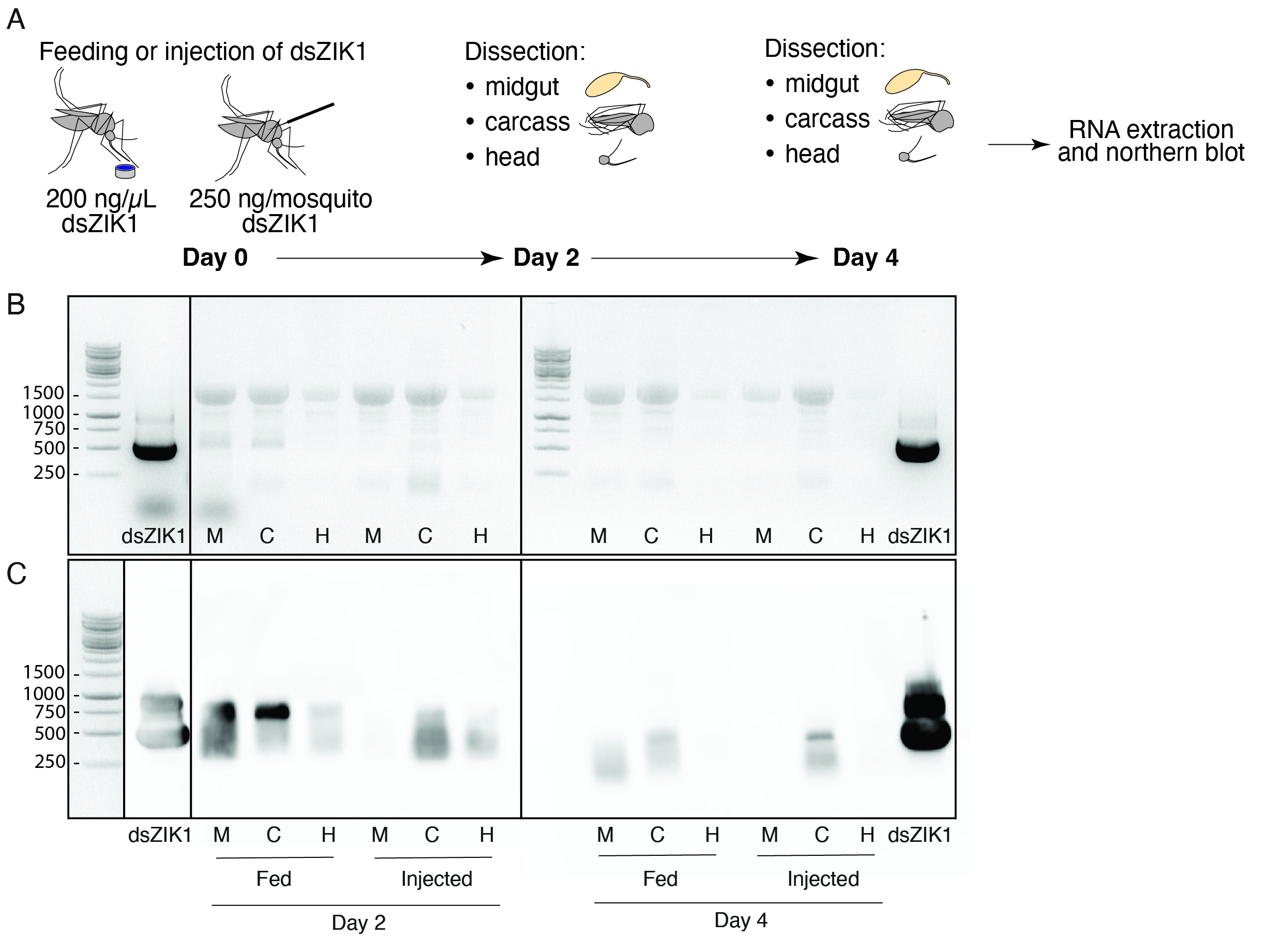
